## Supplementary figures and images for "A *Vibrio cholerae* viral satellite maximizes its spread and inhibits phage by remodeling hijacked phage coat proteins into small capsids"

### Figure 1 Figure Supplement 1

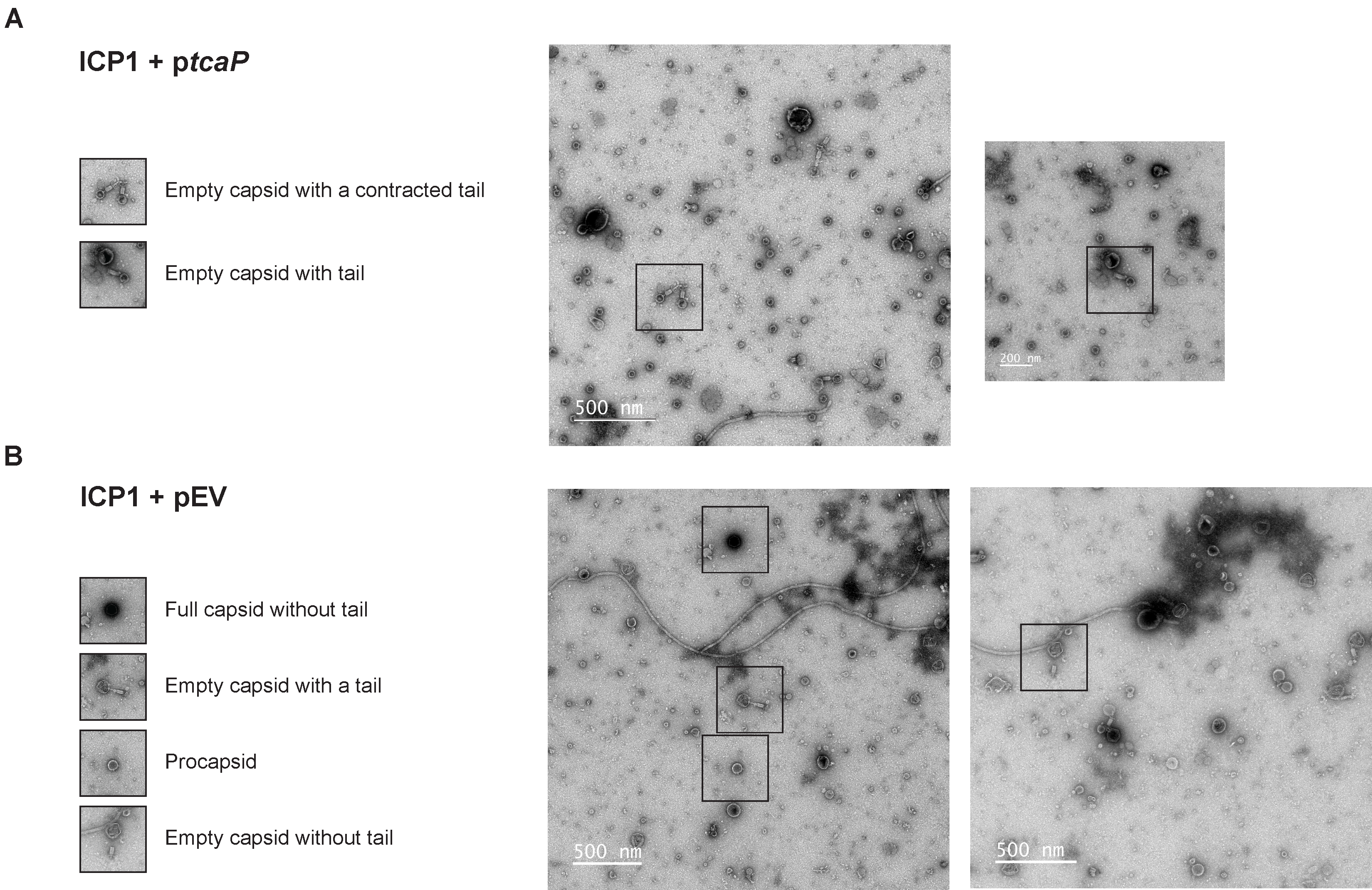

### Figure 3 Figure Supplement 1

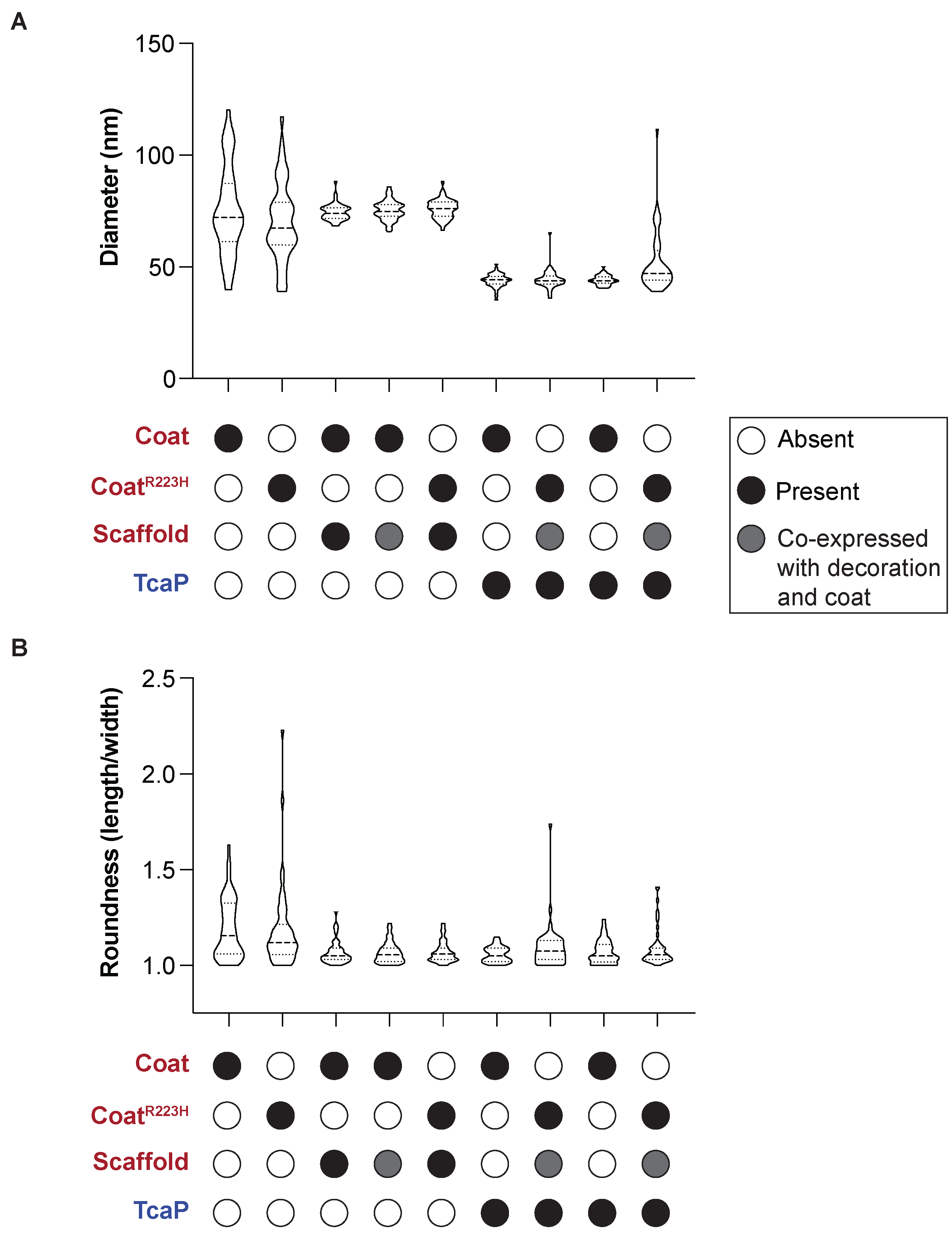

### Figure 3 Figure Supplement 2

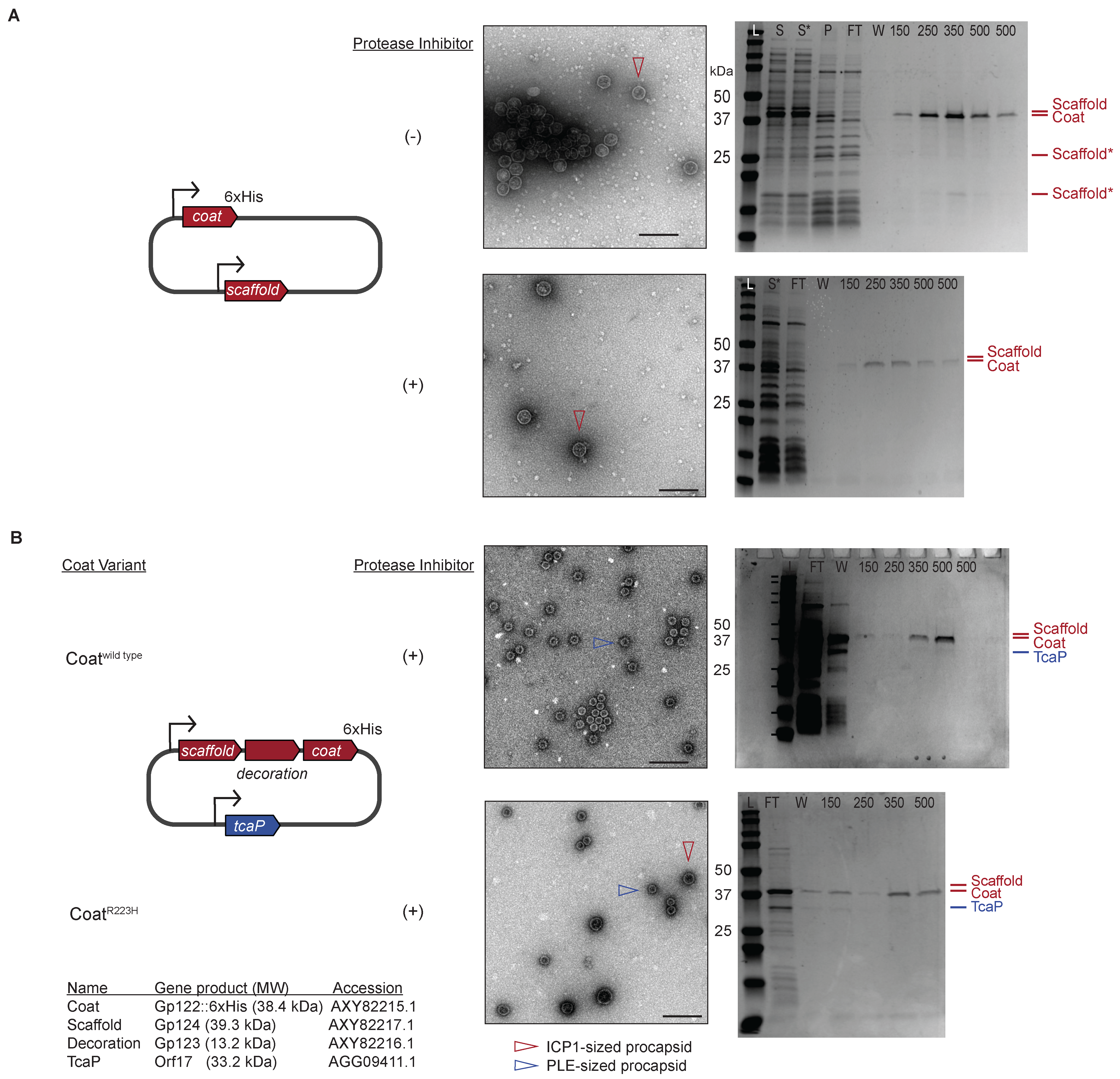

### Figure 3 Figure Supplement 3

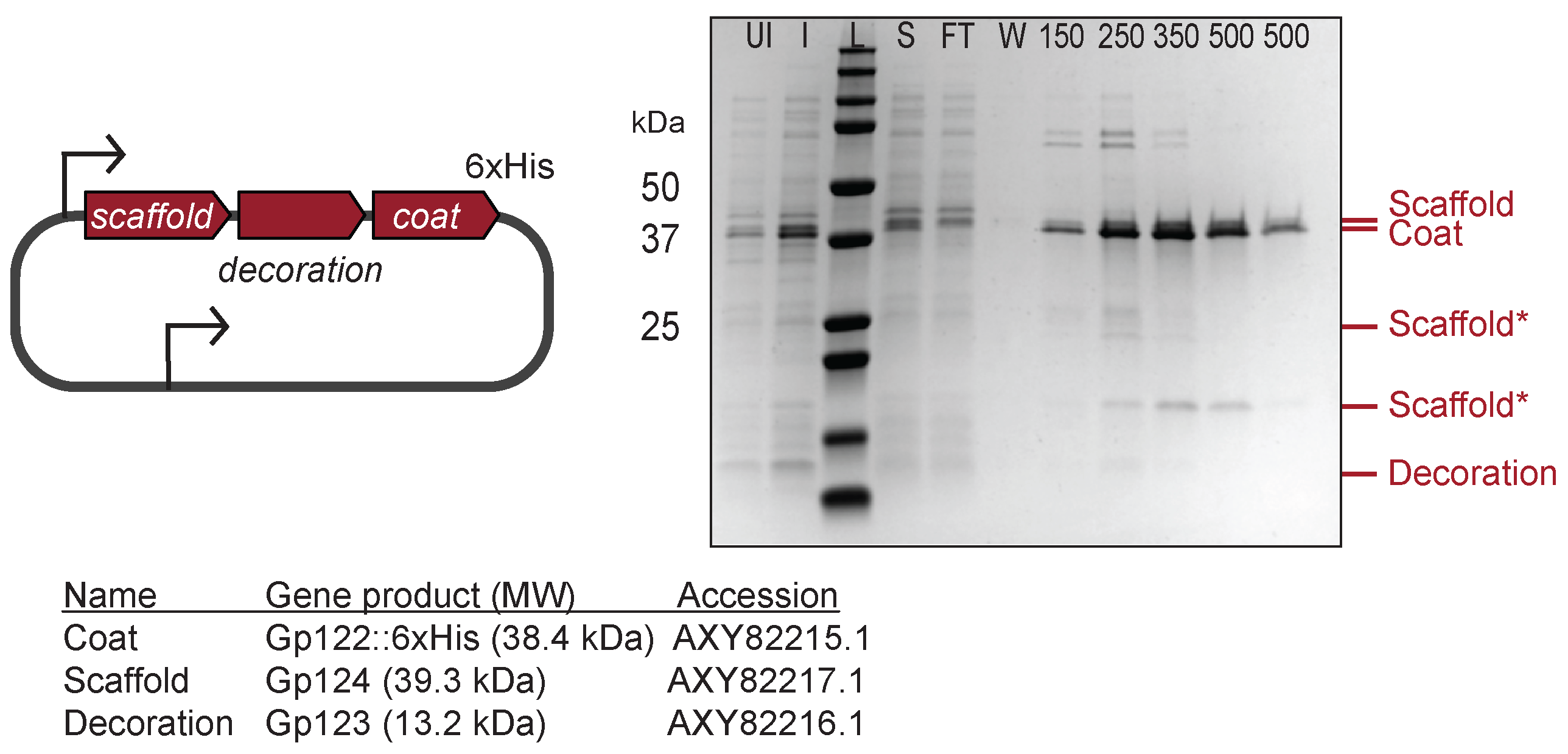

### Figure 3 Figure Supplement 4

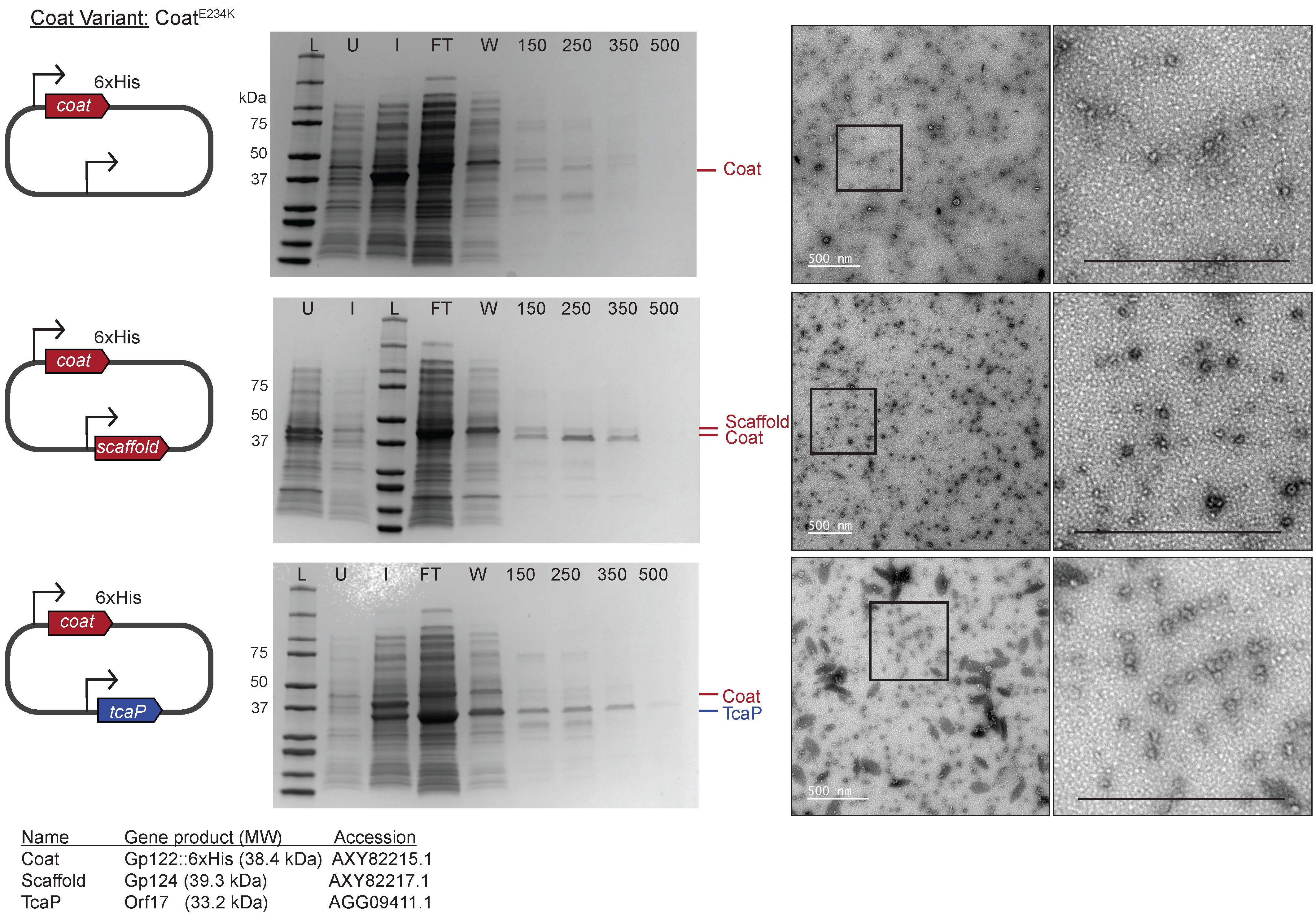

### Figure 4 Figure Supplement 1

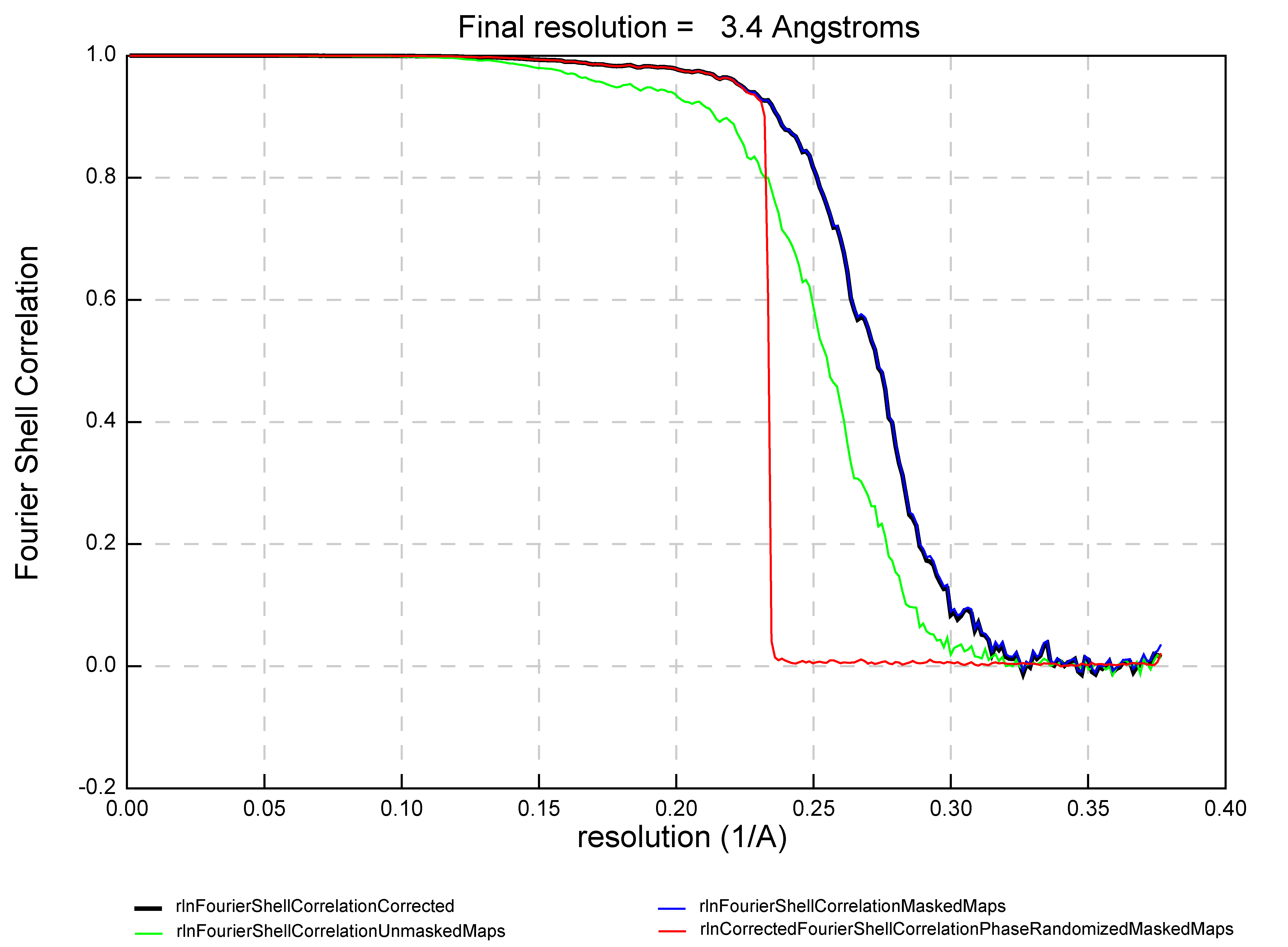

### Figure 4 Figure Supplement 2

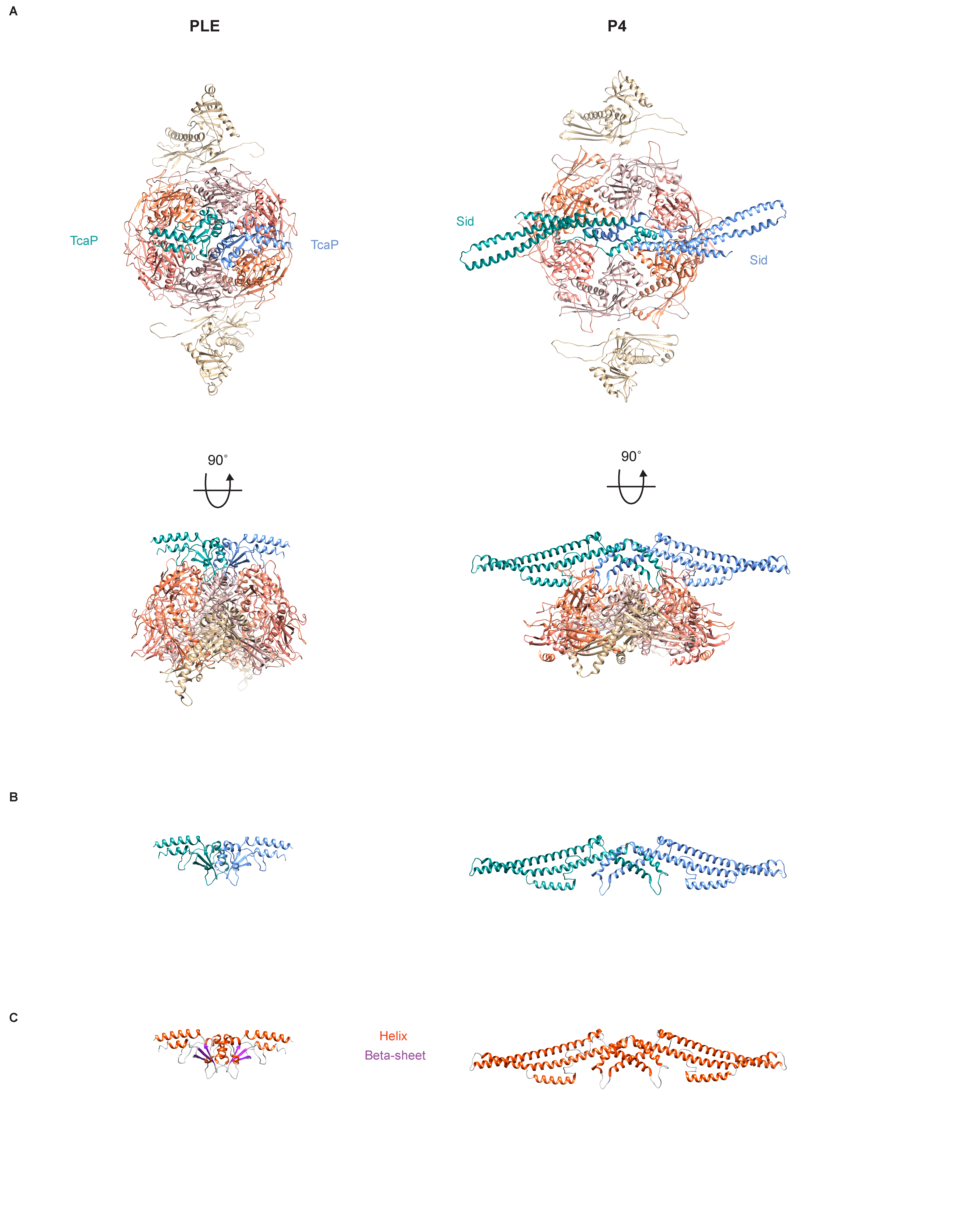

### Figure 5 Figure Supplement 1

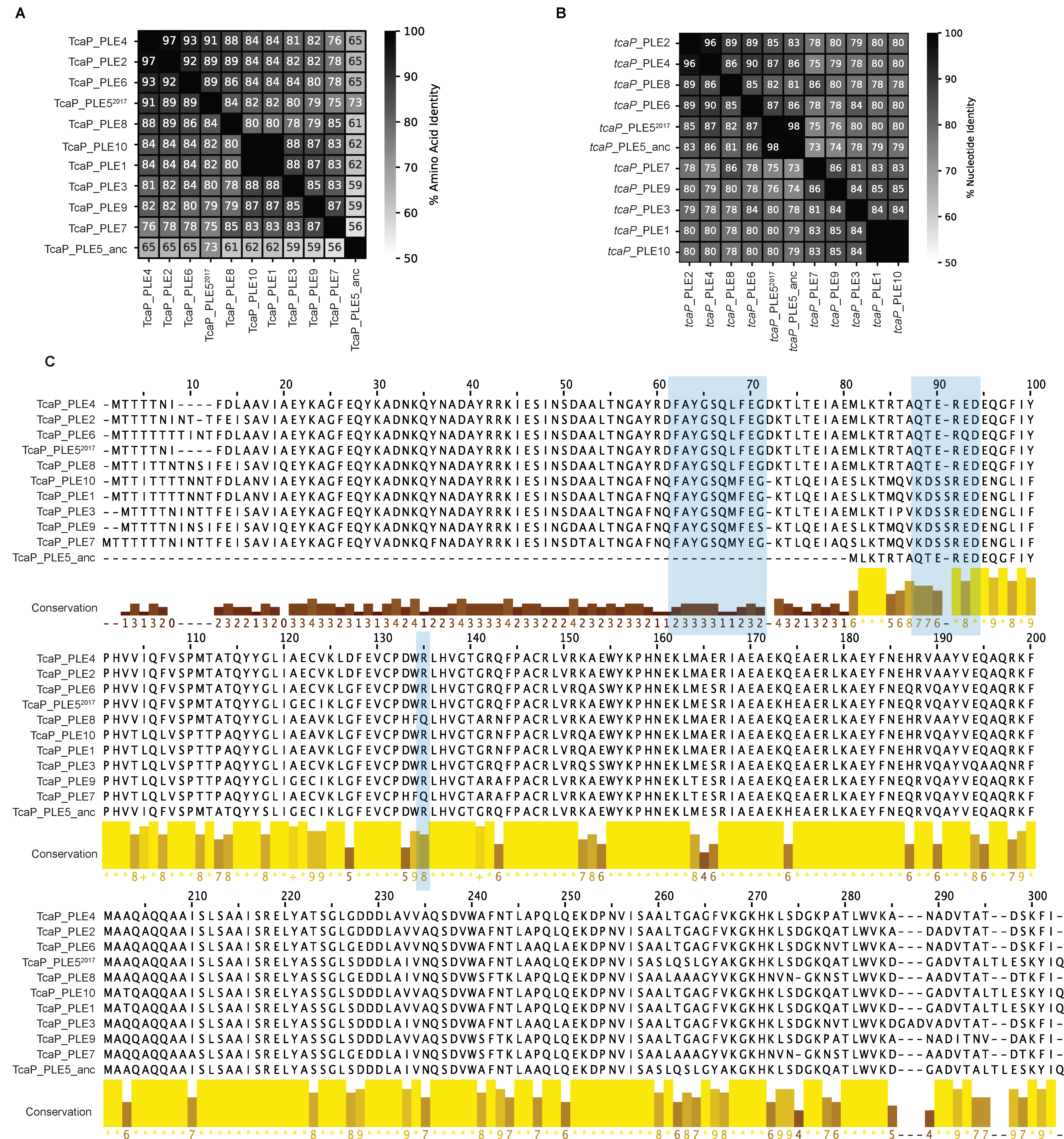
